# Increased extra-axial cerebrospinal fluid and benign external hydrocephalus in early autism

**DOI:** 10.1101/2024.01.04.574160

**Authors:** Gal Ben-Arie, Ilan Shelef, Gal Meiri, Idan Menashe, Ilan Dinstein, Ayelet Arazi

## Abstract

Previous studies have reported that some children with Autism Spectrum Disorder (ASD) below the age of 3-years-old exhibit increased extra-axial cerebrospinal fluid (EA-CSF) volumes, suggesting that transient early CSF circulation abnormalities may characterize a unique etiology in a sub-group of ASD children. Specifically, this sub-group was proposed to exhibit an EA-CSF to total brain volume (TBV) ratio >0.14. It is unknown whether such findings may correspond to an early radiological diagnosis of Benign external hydrocephalus (BEH), which is defined as an enlargement of CSF volumes within the subarachnoid space (SAS) during the first two years of life.

Here we examined a retrospective clinical sample extracted from the National Autism Database of Israel. In 2020, ∼14% of ASD children in the database (n= 102) had completed clinical brain MRI scans at different ages before or after ASD diagnosis. Scans of 83 ASD children fulfilled quality control criteria and were compared to those of 53 age and sex matched typicality developing (TD) children who were referred to the same clinical center for similar clinical reasons. EA-CSF volume and total brain volume (TBV) were quantified from T2-weighted scans using manual labeling and compared across groups.

The results revealed that ASD children scanned before the age of 2-years-old (i.e., before ASD diagnosis) exhibited significantly larger EA-CSF volumes when compared to controls (t(49)=2.89, p=0.006, Cohen’s d=0.82). No significant differences were apparent across groups at older ages. Of those scanned before the age of 2-years-old, ∼16% of the ASD children, and none of the control children, exhibited EA-CSF/TBV ratios >0.14, thereby replicating previous findings from prospective research samples. In addition, retrospective examination of the neuroradiology assessments revealed a BEH prevalence of 33% in the ASD children scanned before the age of 2-years-old, with 80% of these children exhibiting EA-CSF/TBV ratios >0.11.

These findings reveal a clear relationship between quantitative EA-CSF volume analyses and qualitative BEH radiological findings, which characterize a considerable sub-group of ASD children. Accumulating evidence, therefore, suggests that BEH diagnosis in conjunction with a high EA-CSF/TBV ratio by the age of 2-years-old may be indicative of a specific ASD etiology that warrants further research and may offer unique opportunities for intervention.

## Introduction

Autism spectrum disorder (ASD) is a heterogeneous neurodevelopmental disorder characterized by deficits in social communication and the presence of restricted and repetitive behaviors^1^. Over the years numerous theories, mostly based on MRI studies, have suggested that children with ASD may exhibit structural brain differences during early development. Examples include early increases in total brain volume (TBV)^2,3^, abnormal amygdala volumes^4,5^, and cerebellar anatomy^6^, which have all been proposed as potential early biomarkers for ASD. However, many of these findings have not been replicated by studies with larger MRI datasets^7,8^, suggesting that they may not generalize to the broad ASD population. Such mixed findings are common in many areas of ASD research and have led to proposals to abandon the search for a single ASD biomarker^9^. Instead, it may be more useful to search for stratification biomarkers with the potential of identifying ASD subgroups with distinct physiology and etiologies^10^.

One interesting biomarker that may be relevant to a particular sub-group of ASD children is excessive extra-axial cerebrospinal fluid (EA-CSF) in the sub-arachnoid space (SAS) above the frontal lobes^11^. CSF plays a critical role in brain development^12^, ensuring the delivery of growth factors and signaling molecules^13^ while clearing waste and toxins^14,15^. Several studies have reported that a sub-group of young children who go on to receive an ASD diagnosis exhibit increased EA-CSF volumes before the age of 4-years-old^16–18^, suggesting that abnormalities in CSF circulation and/or drainage may be associated with the development of ASD^11^. More specifically, one of these studies reported that 21 of 159 2-4-year-old children with ASD (i.e., 13% of their sample) exhibited an EA-CSF/TBV ratio higher than 0.14^17^. It was suggested that this increased EA-CSF/TBV ratio may indicate a unique etiology of ASD. Two large follow-up studies examining older ASD children did not find any differences in EA-CSF volumes in ASD children 4-years-old and older^19,20^, suggesting that EA-CSF differences are transient and normalize during later childhood.

High EA-CSF/TBV ratios may correspond to a radiological diagnosis of Benign External Hydrocephalus (BEH), which is given to children with abnormally large EA-CSF volumes and otherwise normal neuroimaging findings during the first two years of life^21^. Estimated BEH prevalence ranges from 0.04% ^22,23^ to 0.6% ^24^ of live births. BEH is four times more common in boys than girls (as is the case with ASD) and is usually not treated because EA-CSF volumes return to normal spontaneously without requiring treatment^21^. However, more recent longitudinal follow up studies of BEH cohorts, which included a high percentage (27-30%) of severe BEH cases where shunt operations were performed to reduce intracranial pressure, have reported a relatively high prevalence of psychomotor impairments^25^ and developmental disorders including speech and motor delays, ASD, and ADHD^26^. To the best of our knowledge, previous studies have not examined the correspondence between quantitative measurements of EA-CSF volumes and qualitative BEH diagnoses nor the prevalence of BEH in ASD children.

In the current study we compared EA-CSF and TBV, as manually marked in T2-weighted MRI scans, between children with ASD and age matched controls. Unlike previous studies, we focused on a community-based sample of ASD and typicality developing (TD) children who had been referred for brain MRI scans for various clinical reasons including Seizures, Torticollis, Strabismus, Deafness, Hypotonia, Dysmorphism, and other clinical concerns (Table 2). Importantly, none of the included ASD or TD children had macrocephaly. We hypothesized that younger ASD children, below the age of 4-years-old, would exhibit significantly increased EA-CSF volumes and EA-CSF/TBV ratios compared to controls, and expected to identify a sub-group of young ASD children with EA-CSF/TBV ratios above 0.14, as reported previously by others^17^. We also expected to find a considerable number of ASD children with an early diagnosis of BEH among those with high EA-CSF/TBV ratios before the age of 2-years-old, given the overlapping definition of BEH.

## Materials and methods

### Participants

MRI scans of 102 ASD children and 53 typicality developing (TD) children were retrospectively extracted from the Soroka University Medical Center (SUMC) patient records (Table 1). ASD participants included all children who had been diagnosed with ASD at SUMC between 2013 and 2020 and whose data was available through the National Autism Database of Israel^27,28^. Of the 731 ASD children available in the database at the time, 102 children (∼14%), had been referred to a brain MRI scan at the SUMC radiology department between 2011 and 2020. SUMC is the only clinical center where children insured by the Clalit HMO (who cover 70% of the population in southern Israel) can receive an ASD diagnosis and/or a brain MRI scan, thereby yielding a representative community sample of this geographical area.

**Table 1:** Characteristics of ASD and TD children in the final sample.

| | ASD (mean ± SD) | TD (mean ± SD) |
| --- | --- | --- |
| N | 83 | 53 |
| Age (months) | 36.6 ± 21.5 | 36.4 ± 22.4 |
| Age of ASD diagnosis (months) | 37.7 ± 18.1 | N/A |
| Sex | Females: 24 (29%) | Females: 18 (33.9%) |
| ADOS (N=60, 72%): |  |  |
| Total score | 18.3 ± 6.9 | N/A |
| SA | 14.8 ± 6.3 | N/A |
| RRB | 3.5 ± 1.8 | N/A |
| Calibrated severity score | 7.6 ± 2.7 | N/A |
| Cognitive score (N=38, 46%) |  |  |
| Total score | 66.9 ± 12.3 | N/A |

ASD children were referred to the MRI scan for different reasons (Table 2) either before or after their ASD diagnosis. TD children were selected to match the age range of the ASD children (i.e., 5-100 months old). TD children were referred to MRI for similar reasons (Table 2) and selected if their brain MRI scans showed no radiological findings and their clinical follow-up outcomes were normal (i.e., no evidence of developmental delays). ASD children were excluded from analyses using the following criteria: (1) poor quality of scan (based on visual inspection), (2) missing data (e.g., no T2-weighted scan), and (3) children with microcephaly or macrocephaly (as evaluated by an expert radiologist). This led to the exclusion of 3 ASD children with macrocephaly and 18 ASD children with missing data or poor scan quality, yielding a final ASD sample of 83 children. TD children were selected such that none of these exclusion criteria were met. Sixteen of the 83 ASD children had more than one MRI scan (range: 2-6 scans), of which we included only the earliest usable scan (i.e., each data point corresponded to one individual child).

**Table 2:** Reasons for referral of ASD and TD children to brain MRI.

| ASD | TD |
| --- | --- |
| • Dysmorphism<br>• Strabismus<br>• Hypotonia<br>• Seizures<br>• Deafness<br>• Torticollis<br>• Suspected genetic disorder (including: X-ALD, CHARGE syndrome, Fragile X)<br>• Suspected metabolic disease | • Dysmorphism<br>• Strabismus<br>• Hypotonia<br>• Seizures<br>• Deafness<br>• Torticollis<br>• Suspected genetic disorder (including X-ALD, Neurofibromatosis)<br>• Hearing disability<br>• Suspected endocrine syndrome |
| • Growth hormone deficiency<br>• Premature puberty<br>• Subarachnoid cyst<br>• Setting-sun eye phenomenon<br>• Abnormal mass above the eye<br>• Ocular albinism<br>• Preterm birth<br>• Autism Spectrum Disorder<br>• Developmental delay (including language, motor, and global delay). | • Headaches<br>• Sleep apnea<br>• pilonidal sinus<br>• eye ptosis<br>• Suspected brain lesion per CT<br>• Papilledema<br>• Nasal bridge lesion<br>• Hypophysis abnormalities<br>• Spastic paraplegia |

### ASD diagnosis

As required by Israeli clinical norms, ASD children were diagnosed by both a physician (child neurologist or psychiatrist) and a developmental psychologist, according to DSM-5 criteria^1^. Of the 83 children with ASD included in the final analysis, 60 children also completed the Autism Diagnostic Observation Schedule, 2^nd^ edition (ADOS-2)^29^. ADOS-2 assessments were performed by a clinician with research reliability (Table 1). In addition, 39 of the ASD children also completed a cognitive assessment using the Bayley scales of infant development^30^ or the Wechsler Preschool and Primary Scale of Intelligence (WPPSI)^31^. Cognitive assessments were conducted by a licensed developmental psychologist (Table 1).

### MRI acquisition

All subjects completed MRI scanning with a 1.5T Achieva or a 3.0T Ingenia Philips MRI scanner (Philips Medical Systems, Best, The Netherlands). The 1.5T system was equipped with a 6-channel head coil and the 3T with a 15-channel head coil. On the 1.5T system brain tissue, ventricle volumes, CSF spaces and potential optic nerve (ON) deformity were imaged with a T2-weighted (T2w) turbo spin-echo (TSE) sequence with TE/TR=5730/110 ms, a field-of-view of 210mm, slice width of 4.5mm, in-plane resolution of 0.56 x 0.69m, and a flip angle of 90°. On the 3T system we used a T2-weighted multi-vane turbo spin-echo with TR/TE of 4000/108ms, SPIR fat suppression, slice thickness/gap of 3.8/1.0mm, a field-of-view of 230mm, and a SENSE parallel imaging factor of 1.5. All children in both groups were sedated during the MRI scans. All scans were interpreted clinically by a neuroradiologist with over 15 years of experience (author I.S.).

### MRI analyses

EA-CSF was manually identified on the T2-weighted scans (Fig. 1) in a semi-automated manner by three annotators (A.A and two second year radiology interns) who were blind to the child’s diagnosis. First, a 3D rectangle was manually marked across slices in each of the lateral ventricles. The mean image intensity of the selected voxels was computed and 50% of this value was set as a CSF detection threshold. Second, we selected all voxels with values above the threshold as CSF (note that in T-2 weighted scans, voxels containing CSF have high image intensity values). This procedure was performed repeatedly while altering the threshold and visually inspecting the scans for each participant separately such that the average threshold was set to 54.1% (±6.3%). After selecting the optimal threshold for the participant, annotators used the itk-SNAP software (www.itksnap.org), to visually inspect the CSF labeling and manually correct it. Ventricles were manually removed and only extra-axial CSF voxels above the anterior-posterior commissure horizontal plane were included in the final analysis as also performed by previous studies^16,18–20^. Total brain volumes were estimated by two annotators (Radiology interns) who were blind to the children’s diagnoses (Fig. 1). These estimates were performed using the same T2-weighted scans using the ITK-SNAP software to manually mark all voxels containing the cerebrum (i.e., excluding brain stem and cerebellum) on each slice.

**Figure 1:**
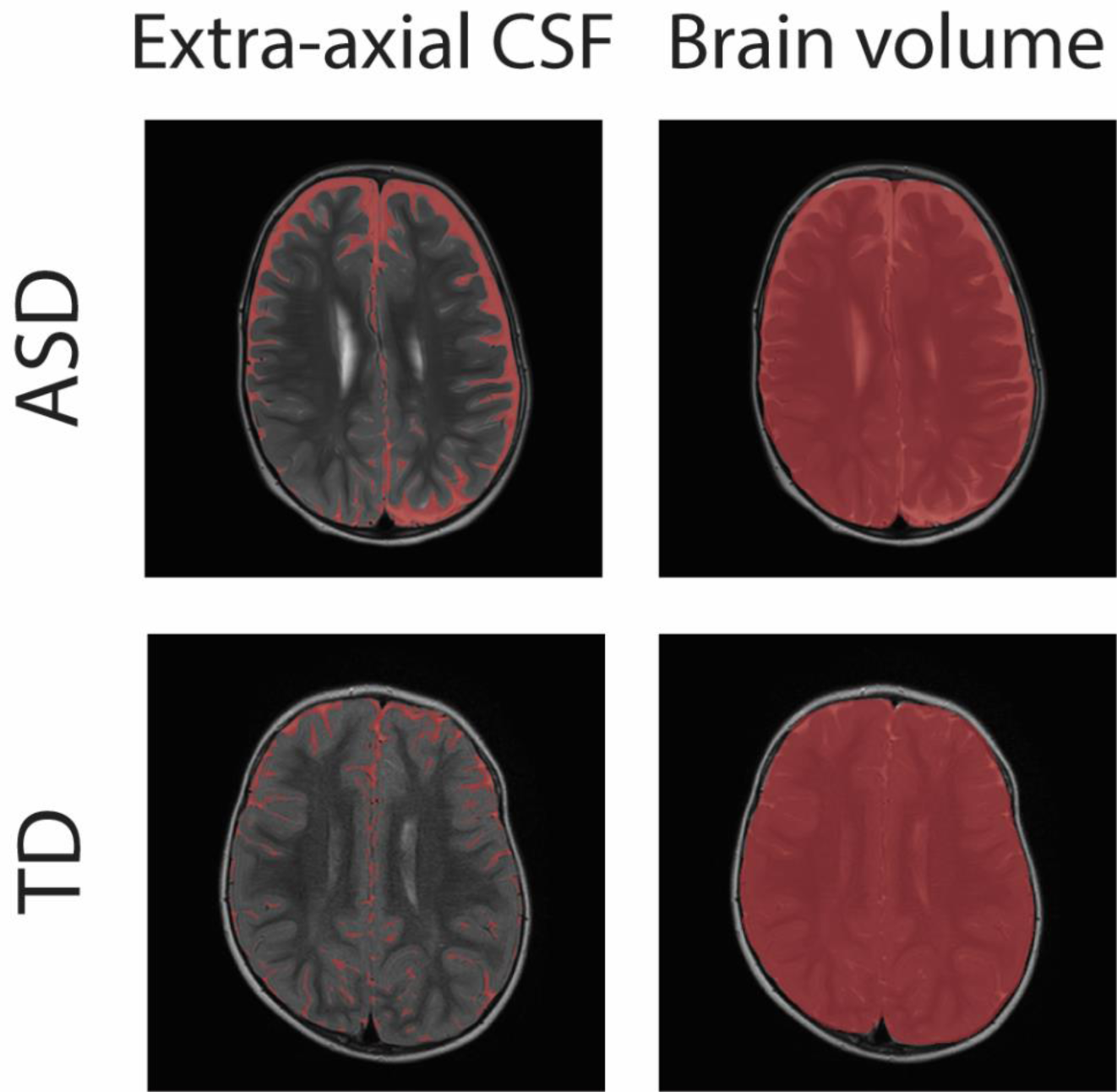
Extra-axial CSF and brain volume labeling. Examples of T2-weighted images with segmentation of extra-axial CSF volumes (left) and brain volume (right) of an ASD child (top) and a TD child (bottom), both scanned at the age of 26 months.

### Statistical analysis

All statistical analyses were performed using MATLAB (Mathworks Inc. United States). Extra-axial CSF volume, brain volume, and their ratio were compared across ASD and TD groups using two-tailed t-tests with unequal variance, and effect size for each comparison was computed using Cohen’s d. Relationships between volumes and age were quantified using Pearson’s correlation coefficients. We also compared findings separately in children of three age groups: 0-24 months (N(ASD)=30, N(control)=21), 24-48 months (N(ASD)=36, N(control)=21) and 49-99 months (N(ASD)=17, N(control)=11). Measures were compared using 2-way ANOVAs with diagnostic group as one factor and age group as a second factor. Follow up comparisons between pairs of age groups were performed using two-tailed t-tests, when the initial results indicated significant differences. We used Bonferroni correction to control for multiple comparisons with the three age groups, adjusting significance thresholds from a p-value of 0.05 to 0.0167.

## Results

ASD children exhibited significantly larger extra-axial CSF volumes compared to age-matched controls (Fig. 2A, left panel; t(134)=3.1, p=0.002, Cohen’s d=0.55). There was a non-significant trend for a negative correlation between extra-axial CSF volume and age in the ASD group (r(83)=-0.18, p=0.1), and no correlation in the control group (r(53)=0.03, p=0.89; Fig. 2A, middle panel). We split the children into three age groups (0-24, 25-48, and >48 months-old) to assess differences across early and late developmental periods. A two-way ANOVA analysis (with age group as one factor, and diagnosis as a second factor) revealed that extra-axial CSF differed significantly by diagnosis (F(1, 130)=8.93, p=0.003), with a marginally significant effect for age (F(2, 130)=2.81, p=0.06), and no significant interaction (F(2, 130)=1.1, p=0.33; Fig. 2A, right panel). Post-hoc analyses performed separately for each age group revealed significantly larger extra-axial CSF in the ASD group only in the youngest age group (t(49)=2.89, p=0.006, Cohen’s d=0.82) and not in the older age groups (age 24-48: t(51)=1.44, p=0.15, Cohen’s d=0.4; age 49-99: t(26)=1.07, p=0.29, Cohen’s d=0.42).

**Figure 2:**
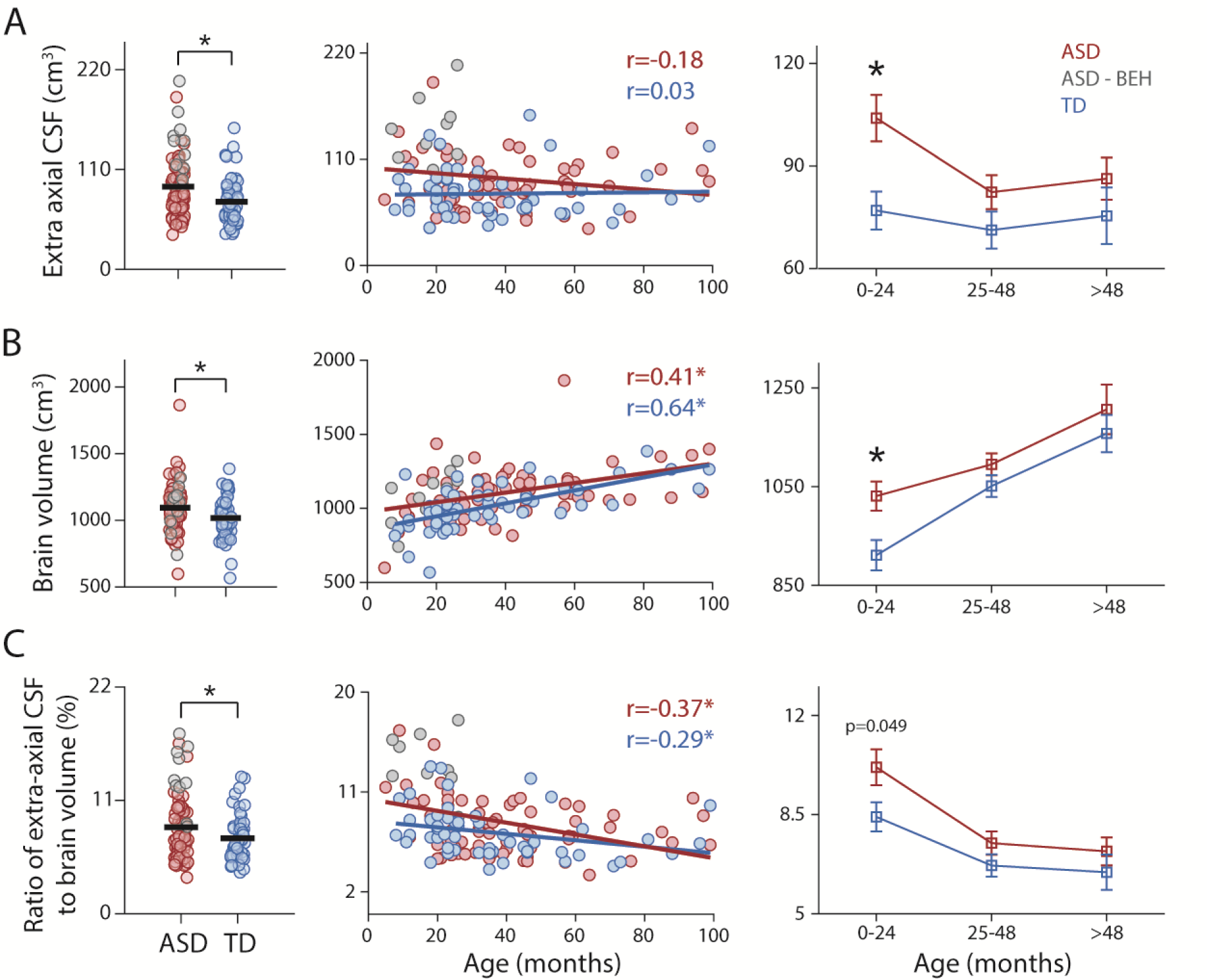
Comparison of extra-axial CSF volume (A), brain volume (B), and their ratio (C) across ASD children without a diagnosis of BEH (red), ASD children with BEH findings (gray), and TD children (blue). Left panels: Comparison across groups (all children). Each circle represents a single child, and the black lines represent group means. Asterisks: significant difference, two-tailed t-test, p<0.05. Middle panels: Scatter plot of each measure and its relationship with age. Lines represent the least squares linear fit and Pearson’s correlation coefficients are noted. Asterisks: significant correlation, p<0.05. Right panels: Comparison across ASD and TD groups, separately for three age groups. Asterisks: significant difference, two-tailed t-test, p<0.0167 (Bonferroni corrected).

Brain volumes were also significantly larger in the ASD group (t(133)=2.65, p=0.009, Cohen’s d=0.47; Fig. 2B, left panel). Brain volumes were significantly positively correlated with age in both groups (ASD: r(82)=0.41, p<0.0001; control: r(53)=0.64, p<0.0001; Fig. 2B, middle panel). A two-way ANOVA analysis revealed that brain volume differed significantly by diagnosis (F(1, 130)=6.76, p=0.01), and across age groups (F(2, 130)=18.29, p<0.0001), with no significant interaction (F(2, 130)=0.97, p=0.38; Fig. 2B, left panel). Post-hoc analyses performed separately for each age group revealed significantly larger brain volume in the ASD group only in the youngest age group (t(49)=2.75, p=0.008).

Finally, we computed the ratio between extra-axial CSF and brain volumes, to determine whether CSF differences across groups were apparent above and beyond differences in brain volume. Ratios were significantly larger in the ASD group (t(133)=2.2, p=0.03, Cohen’s d=0.39; Fig. 2C, left) and were significantly negatively correlated with age in both the ASD (r(82)=-0.37, p<0.0001) and TD (r(53)=-0.29, p=0.035) groups. A two-way ANOVA analysis showed a significant effect for both diagnosis (F(1, 130)=5.34, p=0.02) and age group (F(2, 130)=12.29, p<0.0001) with no significant interaction (F(2, 130)=0.56, p=0.57). Post-hoc analyses performed separately for each age group revealed marginally significant larger ratios in the ASD group only in the youngest age group (t(49)=2, p=0.049, Cohen’s d=0.57, not significant after Bonferroni correction). Taken together, these findings demonstrate an increase in extra-axial CSF volumes that is above and beyond observed increases in brain volume in ASD children under the age of 2-years-old.

### BEH in the ASD group

Of the 83 ASD children examined in this study, 10 children had BEH findings noted in their radiological examinations (gray circles, Fig. 2C). All these children were in the youngest age group (i.e., under 2-years-old) and 8 of the children had an EA-CSF/TBV ratio > 0.11, suggesting a strong relationship between this qualitative radiological finding and the corresponding quantitative measure.

### Reanalysis while excluding ASD children with major radiological findings

ASD and TD children examined in the current study were referred to a clinical MRI scan to assess potential pathologies in brain development. While the included/selected TD children did not have any major radiological findings, some of the ASD children did. We, therefore, performed a second analysis while excluding 14 ASD children who had radiological findings including cortical dysplasia, periventricular leukomalacia, mesial temporal sclerosis, congenital Cytomegalovirus, and posterior fossa cyst that may be expected to impact brain parenchymal or CSF volumes. This analysis, performed with 68 ASD and 53 TD children, reproduced similar results with a marginally significant larger ratio of EA-CSF/TBV in the ASD group (t(118), p=0.09, Cohen’s d=0.31, Fig. 3A).

**Figure 3:**
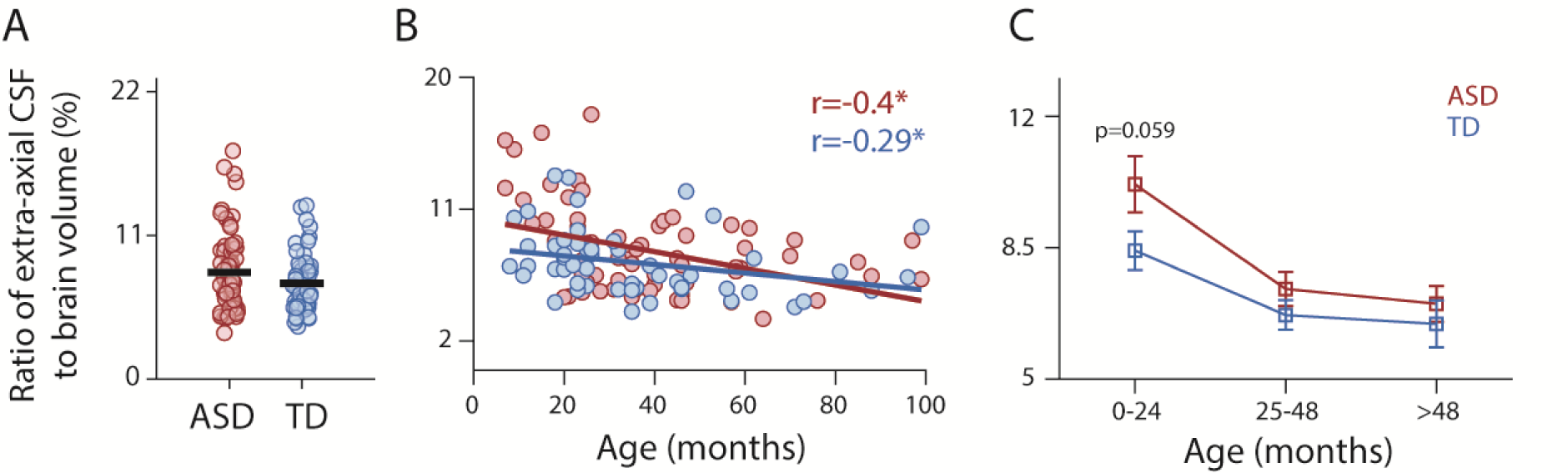
Comparison of extra-axial CSF volume and brain volume ratio across ASD (red) and TD (blue) children after excluding 14 children with radiological findings. (A) Comparison across groups (all children). Each circle represents a single child, and the black lines represent group means. (B) Scatter plot of each measure and its relationship with age. Lines represent the least squares linear fit and Pearson’s correlation coefficients are noted. Asterisks: significant correlation, p<0.05. (C) Comparison across ASD and TD groups, separately for three age groups.

A two-way ANOVA revealed a significant effect for diagnosis (F(1, 130)=4.44, p=0.04) and age group (F(2, 130)=12.42, p<0.0001) with no significant interaction between the two (F(2, 130)=0.7, p=0.5). Two-tailed t-tests performed separately for each age group revealed a marginally significant increase of extra-axial CSF to brain volume ratio in the ASD group only in the youngest age group (t(40)=1.94, p=0.059, Cohen’s d=0.58, Fig. 3C).

### Subgroup of young ASD children with BEH and large EA-CSF volumes

The group differences described above in children under the age of 2-years-old were driven by a subset of children who had extreme EA-CSF volumes. Previous studies have suggested that ASD children with an EA-CSF/TBV ratio ≥0.14 may represent a subgroup with unique pathophysiology^17^. In our study, 5 of 30 ASD children who were younger than 24 months (i.e., 16.7% of this age group) exhibited a ratio ≥ 0.14. In contrast, none of the 21 TD children in this age group exhibited a ratio ≥0.14.

Moreover, 10 of the 30 ASD children in the youngest age group were noted to have BEH findings according to their neuroradiology assessment (gray circles, Fig. 2C). These included 4 ASD children with an EA-CSF/TBV ratio ≥ 0.14 and 4 additional children with a ratio > 0.11. While BEH is a qualitative diagnosis that is based on visual interpretation of MRI scans, these results demonstrate that there is a strong correspondence between BEH and large EA-CSF/TBV ratios. Note that the estimated prevalence of BEH in the general population ranges from 0.04% ^22,23^ to 0.6% ^24^.

### Increased extra-axial CSF may be weakly associated with the severity of ASD symptoms

We estimated the relationship between the severity of autism symptoms, as assessed using the ADOS-2 CSS (see methods), and each of the anatomical measures. Of the 83 children included in the main analyses, 60 children completed the ADOS assessment. We found marginally significant correlations between their ADOS-2 CSS scores and EA-CSF volumes (r(60)=0.23, p=0.07) as well as EA-CSF/TBV ratios (r(60)=0.23, p=0.08). There was no significant correlation with TBV (r(60)=0.01, p=0.97).

## Discussion

Accumulating evidence suggests that excessive EA-CSF volumes during early development characterize a potentially unique sub-group of ASD children. In the current study large EA-CSF volumes, apparent above and beyond increases in TBV, with EA-CSF/TBV ratios exceeding 0.14, were apparent in ∼16% of ASD children scanned before the age of 2-years-old (Fig. 2). This finding corresponds well with a recent report where ∼13.2% of 3-year-old ASD children exhibited the same ratio ^17^. Moreover, two previous studies reported that early EA-CSF volumes at 6 and 24 months of age were significantly larger in high-risk baby siblings of ASD children who also developed ASD versus those who did not^16,18^. Hence, similar findings were apparent in the current retrospective analysis of children scanned for clinical reasons in a medical center setting and multiple prospective studies performed in a research university setting.

ASD children scanned at older ages in the current study (up to 8.3 years old) did not exhibit significant EA-CSF or TBV enlargements relative to controls (Fig. 2). Indeed, two large studies have recently reported that EA-CSF volumes of ASD children above the age of four do not differ significantly from those of controls^19,20^. Taken together, these findings suggest that excessive EA-CSF volumes and EA-CSF/TBV ratios in the sub-group described above are a transient characteristic that appears during early development and disappears later in childhood as the children mature.

This transient phenomenon corresponds well with a diagnosis of BEH, which is defined qualitatively as an abnormal increase in CSF volume within the SAS that appears during the first two years of life and usually disappears spontaneously with time^21^. Remarkably, one third of the ASD children scanned before the age of 2-years-old in the current study had BEH findings according to the examining neuroradiologist. Importantly, 4 of these children had EA-CSF/TBV ratios >0.14 and an additional 4 children had ratios > 0.11 (Fig. 2), thereby establishing a clear correspondence between this qualitative diagnosis and quantitative measure.

The fact that a considerable number of children who were later diagnosed with ASD exhibited BEH findings and high EA-CSF/TBV ratios before the age of 2-years-old, suggests that BEH may not be a benign finding, and that early CSF circulation abnormalities may play a role in the ASD etiology of a specific sub-group of children^11^.

### ASD etiology involving impaired EA-CSF circulation?

Recent studies have revealed that CSF circulation plays many critical roles during early brain development^32^ including delivery of growth factors and essential metabolites^13^ as well as removal of waste^14^. Indeed, multiple types of CSF circulation impairments including congenital and acquired forms of hydrocephalus can cause severe developmental disorders^33,34^ with underlying mechanisms that are poorly understood^35^. The findings reported in this and previous studies^16,17^ suggest that a milder, transient form of hydrocephalus during early development (BEH) may be associated with later development of ASD. Further exploration of the potential mechanisms underlying this relationship is highly warranted and may yield targeted interventions for this specific sub-group of ASD cases.

### Limitations

The current study had several limitations. First, we compared ASD and TD participants who were referred to a brain MRI scan for clinical reasons and may not faithfully represent the ASD and TD general populations. Note, however, that prospective research samples in university setting also do not necessarily generalize to the broad population due to a variety of common social, racial, and cognitive sampling biases^36,37^. Hence the current sample, with its unique characteristics, extends previous findings from other types of samples. Second, our quantification of EA-CSF and TBV was based on T2-weighted scans with poorer spatial resolution than previously utilized T1-weighted MRI scans, which may yield less accurate annotation of TBV and EA-CSF volumes. Third, the manual annotation of EA-CSF and TBV used in the current study would not be conducive to work with larger datasets where use to automated techniques would be required. While automated techniques have been developed for analyses of T1-weighted scans^18^, extending these techniques to T2-weighted scans has not been carried out to date. Finally, the qualitative definition of BEH suggests that there is likely to be variability in how different radiologists allocate this diagnosis to different children. We did not examine inter-rater reliability across radiologists in the current study.

## Conclusions

Contemporary ASD research highlights the large heterogeneity of behaviors and underlying genetics and biology apparent in different individuals with ASD^38^. This has motivated research focused on the identification of more homogeneous sub-groups of ASD individuals who may share specific etiologies and benefit from particular interventions^39^. The results of the current study further support the hypothesis that excessive EA-CSF volumes during the first two years of life characterize a specific sub-group of children who develop ASD. They suggest a potentially unique etiology involving early transient abnormalities in CSF circulation that resolve spontaneously by the age of 4-years-old, which may impact ∼15% of ASD children. The current study demonstrates the considerable correspondence between quantitative measures of EA-CSF volumes and a qualitative radiological diagnosis of BEH, which was highly prevalent in the current study. Further studies aiming to elucidate the biological mechanisms underlying this clinical phenomenon in children with ASD are highly warranted.

## Data availability

The data are available from the corresponding authors upon reasonable request.

## Funding

This work was supported by an Israel Academy of Sciences Adams Fellowship to A.A., Israel Science Foundation grant 1150/20 to I.D, and Azrieli Foundation grant to I.D, I.M, and G.M.

## Competing interests

The authors report no competing interests.

## Notes

### Competing Interest Statement

The authors have declared no competing interest.

